# Integrated Single-Fiber Multi-Omics Links an Inflammatory-Associated Myofiber State to Altered Myosin Dynamics in Patients with ICU-acquired weakness

**DOI:** 10.64898/2026.02.16.706099

**Authors:** Alexandra M. Winant, Roger Moreno-Justicia, Lucia Paolini, Wout J. Claassen, Coen A.C. Ottenheijm, Atul S. Deshmukh, Stefano Cattaneo, Simone Piva, Nicola Latronico, Robert A.E. Seaborne, Julien Ochala

## Abstract

Skeletal muscle dysfunction is a pervasive complication of critical illness that worsens survival and recovery, yet remains poorly explained by current clinical or molecular markers. To directly connect disease-associated molecular states to the contractile machinery, this study combined sequential functional, transcriptomic, and proteomic profiling of the same single human skeletal myofibers from critically ill patients in the intensive care unit with acquired weakness (ICU-AW) and controls. Despite marked donor-level heterogeneity, integrated analysis revealed a subtle yet conserved myofiber state enriched in ICU-AW, characterized by inflammatory and chemotactic gene programs, intracellular structural remodeling, and bioenergetic adaptation. Nineteen features were significantly altered at both RNA and protein levels from the same myofiber, linking an inflammatory transcriptional landscape to a proteomic shift toward mitochondrial and translational machinery and away from membrane-associated signaling. Functionally, fibers in this state displayed selectively disrupted myosin dynamics, evidenced by prolonged ATP turnover time of myosin heads in their super-relaxed conformation, implicating altered myosin energetics as a contributor to muscle dysfunction. These findings define a discrete, disease-associated myofiber state and establish an integrative single-fiber framework for connecting multi-omic heterogeneity to molecular motor function in complex human disease.

**Graphical Abstract:** Single-fiber multi-omic and functional analysis reveals a stress-adapted myofiber state in ICU-AW. Specifically, for the present study, myofibers from ICU-AW donors and control donors were isolated and functionally profiled for myosin dynamics before being split for simultaneous transcriptomic and proteomic analysis. Integrated analysis then identified a reproducible fiber phenotype enriched in ICU-AW, characterized by inflammatory transcriptional signatures coordinated with mitochondrial proteomic remodeling and altered myosin super-relaxed state energetics.

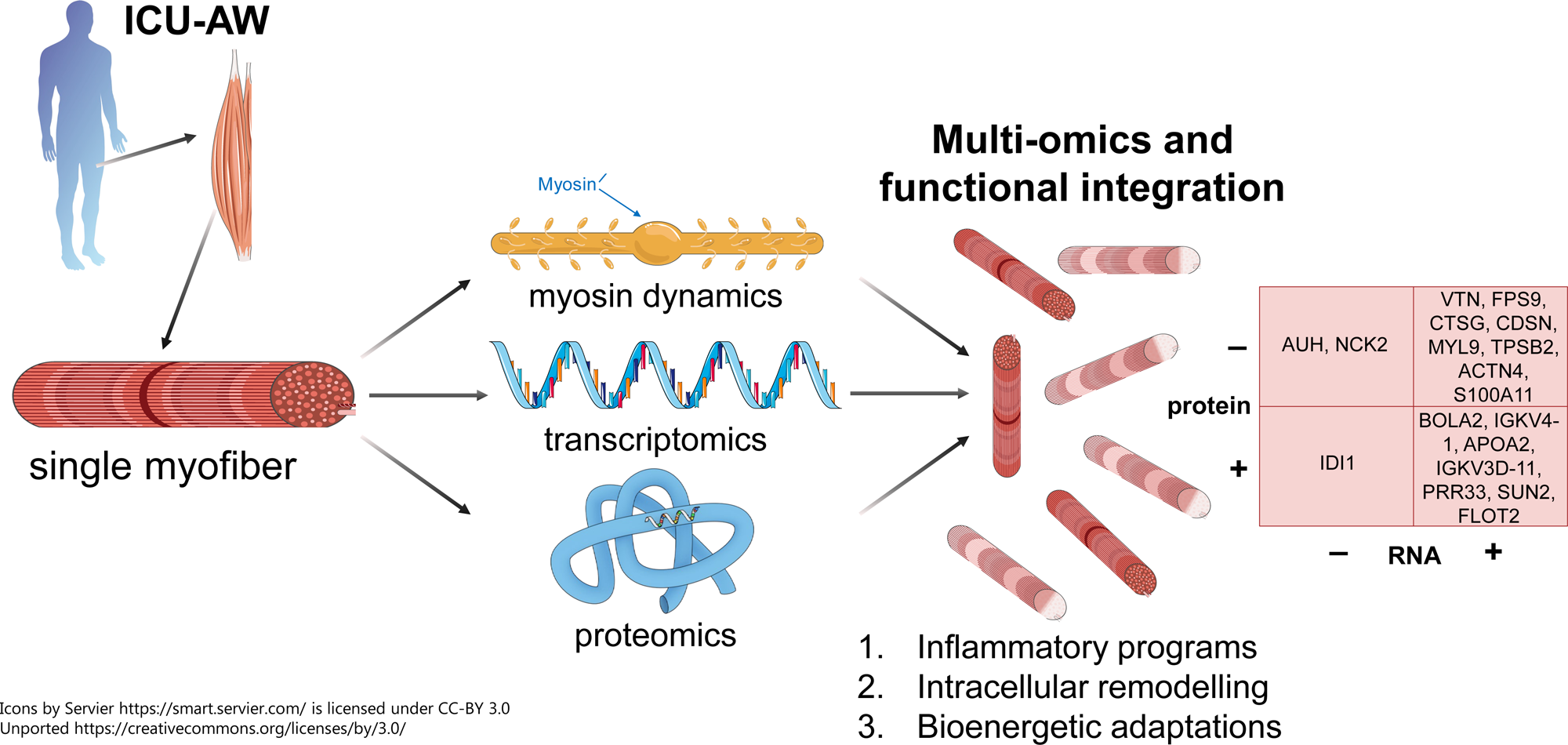

## Introduction

Skeletal muscle is a central determinant of whole-body health, contributing to locomotion, posture, metabolic homeostasis, and endocrine signaling throughout the lifespan. Loss of muscle mass or function can occur rapidly and is associated with impaired mobility, metabolic dysfunction, prolonged hospitalization, and increased mortality across aging, chronic disease, and critical illness (1, 2). Hence, skeletal muscle maintenance is crucial and depends on coordinated regulation of multiple cell types and cell states within a structurally and molecularly complex tissue microenvironment (3).

Myofibers are the most abundant cell type in skeletal muscle and are classically categorized by myosin heavy chain isoform composition and metabolic properties: slow oxidative (type I) and fast glycolytic (type IIA and IIX) fiber types (3). While this classification captures key aspects of contractile specialization, it underestimates the molecular diversity within and between individual myofibers (4). As multinucleated syncytia, myofibers exhibit spatially organized and non-uniform regulation along their length (5). Muscle tissue homeostasis is further shaped by satellite cells, fibroblasts, endothelial cells, and immune cells that collectively support tissue structure and function (3) The resulting cellular and molecular heterogeneity of the muscle niche poses a challenge for achieving high-resolution, cell-specific measurements in human disease (6).

The vulnerability of skeletal muscle is particularly evident in critical illness, where patients may lose nearly 2% of muscle mass per day during the first week of intensive care unit (ICU) admission (7). This rapid, and often profound, ICU-acquired weakness (ICU-AW) is incompletely explained by the admission diagnosis, treatment protocols, or broad measures of muscle atrophy (8). Mechanistic studies implicate increased inflammation and cytokine production, altered calcium homeostasis, enhanced proteolysis, and bioenergetic failure as contributors to ICU-acquired muscle dysfunction (9). There is a preferential loss of myosin and myosin-associated proteins and a consistent reduction in myosin-to-actin ratios (10–15). However, how these processes converge at the level of individual myofibers remains incompletely understood.

Myosin molecules dynamically cycle between biochemical states with distinct energetic properties, including the energy-sparing super-relaxed (SRX) and more active disordered-relaxed (DRX) conformations, which together contribute to basal ATP turnover time and energetic demand in resting muscle (16). Disruption of the balance between myosin states can impair muscle bioenergetics and contractile capacity even in the absence of altered muscle structure, suggesting that altered myosin dynamics might link molecular stress responses to functional weakness during critical illness (16). Yet, how these energetic states are regulated within individual human myofibers in the setting of systemic inflammation, oxidative stress, and mitochondrial dysfunction is largely unexplored (17).

Multiple transcriptomic (11, 12, 18–21) and other high-throughput (22–24) studies have mapped molecular changes in skeletal muscle during ICU-AW, highlighting perturbed signaling pathways and altered gene expression programs. However, transcriptional changes do not consistently translate into protein-level or functional alterations, while bulk analyses obscure cell- and state-specific behaviors of individual myofibers. Prior work has shown that single-myofiber analyses can reveal physiologically relevant diversity in protein expression and function that is substantially reduced in monogenic myopathies, but these approaches typically capture only one molecular layer per fiber (4) Here, building upon our recent work (25), an integrated single-fiber approach was applied that combined myosin functional measurements with transcriptomic and proteomic profiling from the same human myofiber to define molecular states associated with ICU-AW. By linking gene expression, protein abundance, and myosin SRX/DRX dynamics within individual fibers, this study identified a reproducible, disease-enriched myofiber state characterized by inflammatory signaling, intracellular structural remodeling, and bioenergetic adaptation. The presence of this state across ICU-AW patients, supports a fiber-specific molecular and functional program induced by critical illness. Our integrated approach provides a framework for validating molecular signals directly within the cellular and biomechanical context.

## Results

### Workflow integrates single-fiber transcriptomics, proteomics, and myosin dynamics

Originally, we developed a novel workflow enabling sequential isolation and analysis of the same myofiber across multiple molecular and functional layers. Individual myofibers from the quadriceps femoris of eight ICU-AW patients and eight control donors (orthopedic surgery patients) were manually isolated and divided into two segments (Table 1). The first segment underwent Mant-ATP chase experiments to measure myosin ATP turnover kinetics (SRX and DRX states); the second segment was processed for simultaneous transcriptomic and proteomic analysis, with careful registration to enable later dataset integration.

**Table 1.**

| Characteristic | Case<br>N = 8 | Control<br>N = 8 |
| --- | --- | --- |
| Sex, N° (%) |  |  |
| Male | 7 (87.5%) | 4 (50%) |
| Female | 1 (12.5%) | 4 (50%) |
| Age, arithmetic mean | 60 | 64 |
| Reason for admission, N° (%) |  |  |
| Sepsis or septic shock | 2 (25%) | 0 (0%) |
| ARDS | 4 (50%) | 0 (0%) |
| Trauma | 2 (25%) | 8 (100%) |
| Days between ICU Admission and Biopsy |  |  |
| 6 | 1 (12.5%) | - |
| 7 | 3 (37.5%) | - |
| 8 | 2 (25%) | - |
| 12 | 1 (12.5%) | - |
| 13 | 1 (12.5%) | - |

For each donor, 12 myofibers were initially isolated and processed through both functional and multi-omic pipelines. Transcriptomic libraries were prepared with individual molecular barcodes enabling pooled sequencing. Quality control filtering removed transcriptomic samples with gene counts or feature abundance ±3 median absolute deviation (MAD), and proteomic samples with more than 50% missingness (Fig. 1a-b).

**Figure 1.**
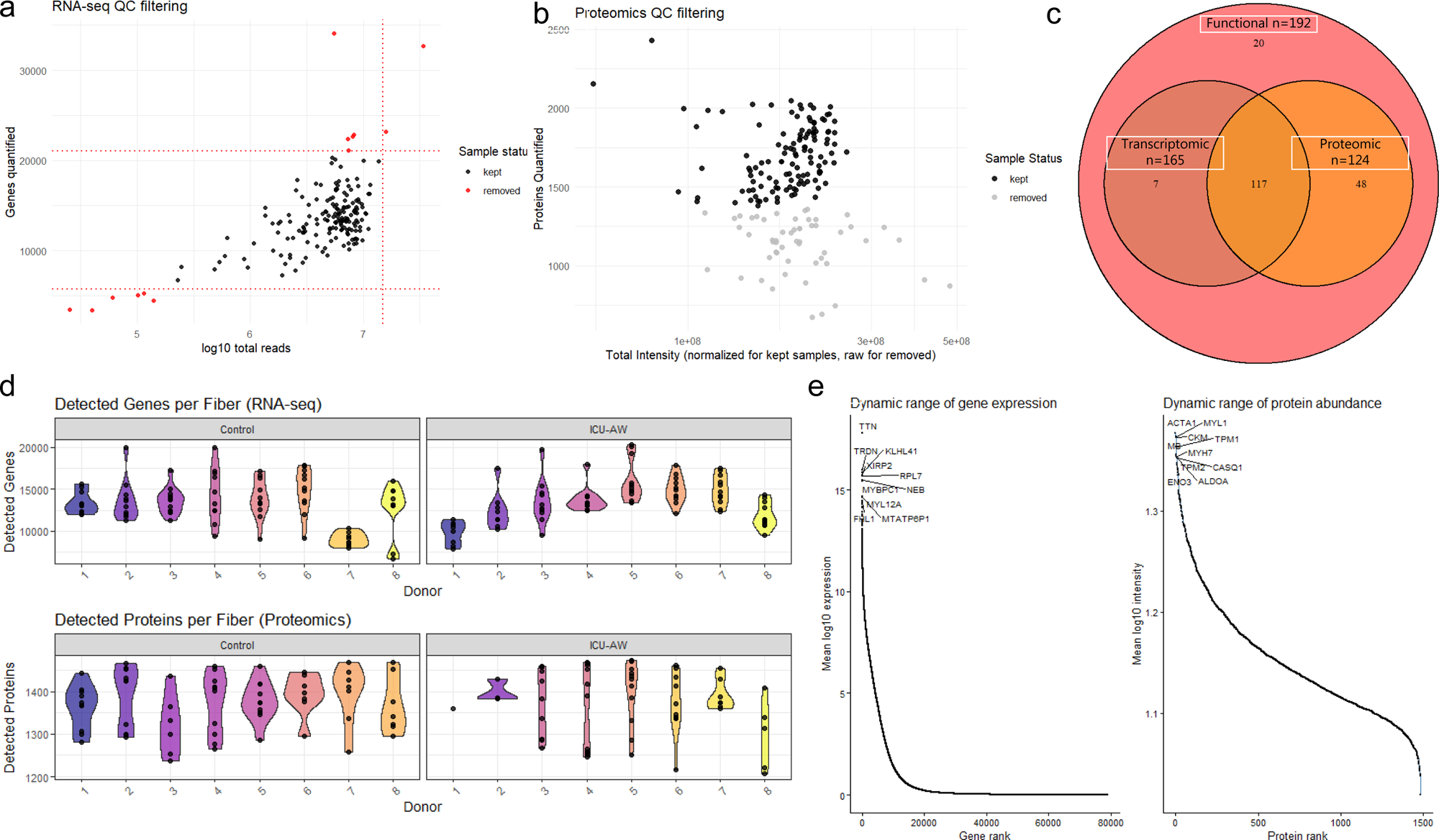
Quality control metrics by dataset. a) Filtering of RNA-seq data. Red dotted lines mark 3 median absolute deviations. Samples outside this cutoff were excluded (red dots), while black dots show retained samples. b) Filtering of proteomic samples. Total intensities were normalized across kept samples (≤50% sample, ≤70% protein missingness). Samples failing filter are plotted in grey using raw intensity to show QC status. c) Overlap of retained samples after filtering. d) Detected features per fiber. Violin plots show detected features per fiber for RNA (top) and protein (bottom). e) Dynamic range of mRNA and protein abundance. Genes (RNA) and proteins were ranked by their mean abundance across fibers following log-transformation.

This resulted in 165 fibers passing transcriptomic quality control (QC) and 124 passing proteomic QC, with 117 fibers common to both datasets (Fig. 1c). Transcriptomic sequencing achieved an average depth of 5.7 × 10⁶ reads per sample with 13,456 ± 2,682 unique genes detected. Proteomic analysis detected 1,372 ± 70 proteins per sample with 8% average missingness, after filtering out proteins not present in at least 70% of samples from the same condition (Fig. 1d). This depth and coverage highlights that our single-fiber, multi-omic approach has sufficient resolution for detecting both common and subtle molecular features in human skeletal muscle samples (Fig. 1e). Workflow and successful number of fibers analyzed per subject are presented in Table S1.

### Subset of ICU-AW myofibers display distinct inflammatory and stress-response gene programs without genome-wide expression shifts

Differential expression analysis was performed on transcriptomic data pseudobulked by donor to account for nested donor structure (donor-level comparison). Although no differentially expressed genes (DEGs; FDR < 0.05) were identified when directly comparing control and ICU-AW groups (Fig. 2a), ranked gene set enrichment analysis (GSEA; Hallmark pathways) revealed distinct higher-level transcriptional signatures between conditions (Fig. 2b). Fibers from ICU-AW patients were characterized by upregulation of stress- and inflammation-related pathways including reactive oxygen species pathway, protein secretion, IL6-JAK-STAT3 signaling, MYC targets, and DNA repair. Conversely, pathways related to metabolism, immune homeostasis, and tissue maintenance (including adipogenesis, myogenesis, and interferon responses) were downregulated in fibers from ICU-AW patients (Fig. S1).

**Figure 2.**
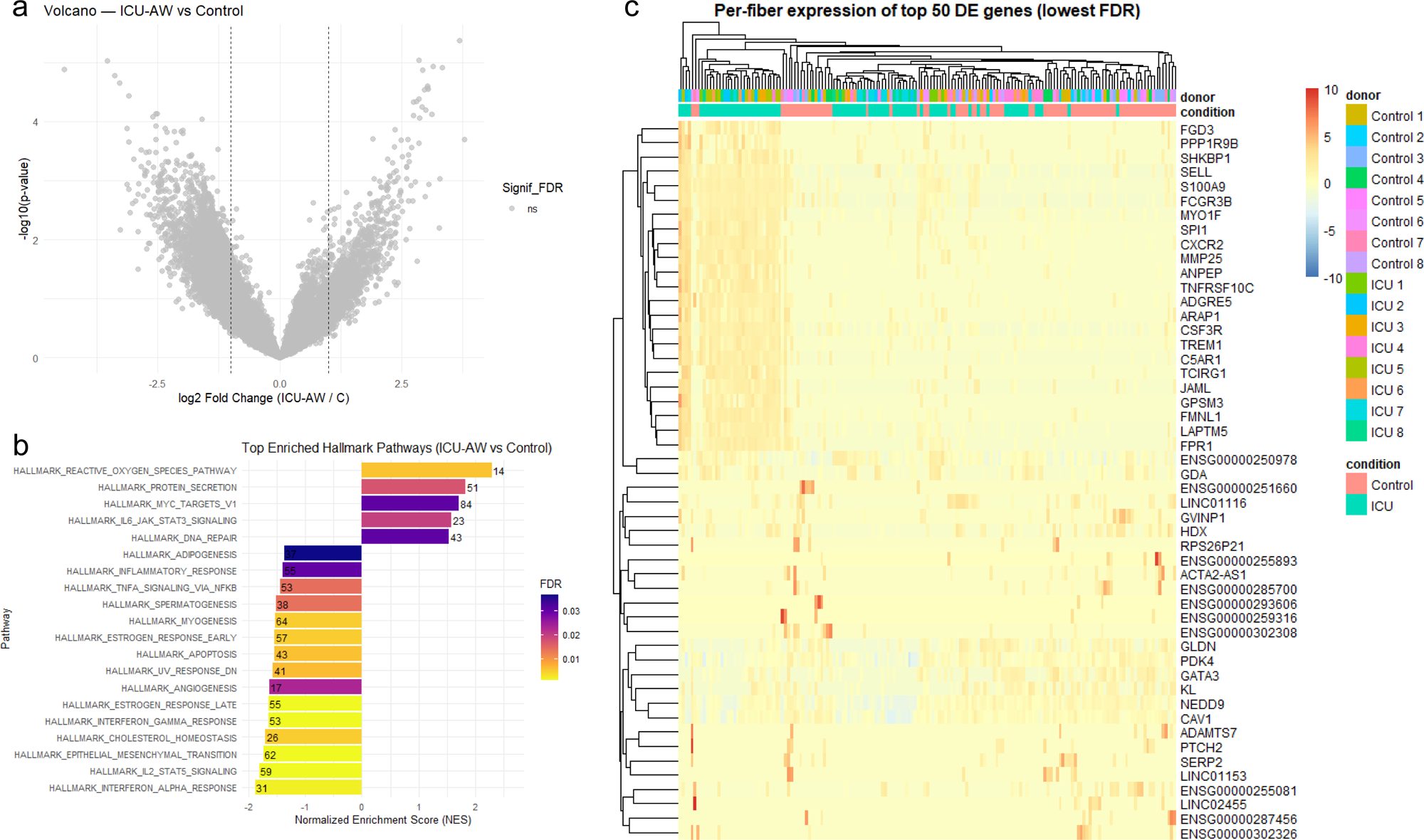
Transcriptional alterations in single myofibers associated with ICU-AW. a) Volcano plot of DEGs between ICU-AW and control samples. Fibers pseudobulked by donor. Gray dots indicate non-significant genes. b) Hallmark pathway enrichment analysis for ICU-AW. The top 20 significantly enriched pathways are ranked by normalized enrichment score (NES) and colored by false discovery rate (FDR). Numbers adjacent to each bar indicate the leading-edge gene count. c) Heatmap of top 50 genes (by lowest FDR) from pseudobulk DE analysis on single fibers.

Hierarchical clustering of the top 50 DEGs (by FDR from donor-level analysis) revealed variation in gene expression patterns across fibers (Fig. 2c). A distinct subcluster emerged with consistently elevated expression of the top 23 DEGs. Strikingly, this subcluster was predominantly composed of myofibers from ICU-AW patients (n = 32) with only one control fiber, suggesting a conserved transcriptional phenotype within the ICU-AW population (Fig. 3a). The subcluster is not defined by fiber type (with similar myosin heavy chain composition to the general dataset) or feature counts (Fig. S2). This subcluster was designated the ‘Inflammatory’ cluster.

**Figure 3.**
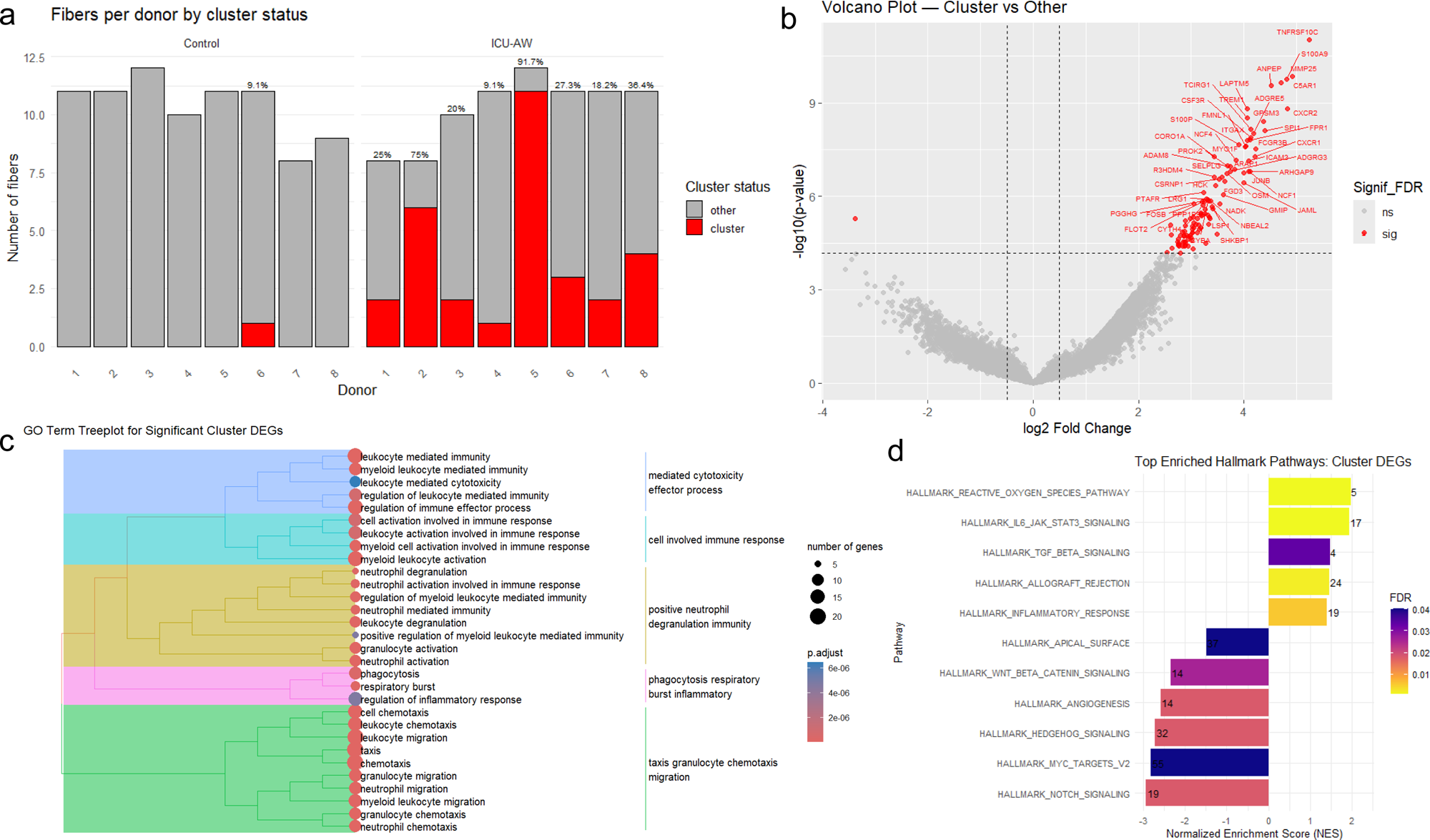
Cluster-Specific Composition and Transcriptional Signatures in Single Myofibers. a) Cluster composition by donor and condition. Each bar shows total number of fibers, with fibers in cluster colored red and quantified in percentage. b) Volcano plot of DEGs between cluster and other. Fibers pseudobulked by donor and cluster status. Red dots indicate significant genes. Top 50 features are labelled. c) Treeplot of top 30 enriched GO terms for cluster DEGs. This visualization shows clustering among biologically related GO categories. d) Hallmark pathway enrichment analysis for cluster DEGs. The significantly enriched pathways are ranked by normalized enrichment score (NES). Bar length represents the NES, and bar color indicates the FDR-adjusted p-value. Numbers adjacent to each bar indicate the leading-edge gene count.

To characterize the Inflammatory cluster transcriptional phenotype, differential expression analysis was performed comparing cluster-assigned versus non-cluster fibers, aggregated by donor (Fig. 3b). This identified 108 DEGs, of which 107 were upregulated in cluster fibers. Gene Ontology (GO) enrichment analysis of identified DEGs revealed strong enrichment for immune-related processes including mediated cytotoxicity, immune cell activation, neutrophil degranulation, respiratory burst, and granulocyte chemotaxis (Fig. 3c). Hallmark pathway analysis confirmed significant upregulation of reactive oxygen species, cytokine signaling, and immune activation pathways, with corresponding downregulation of developmental and proliferative pathways (WNT/β-catenin signaling, angiogenesis, Hedgehog signaling, MYC targets, Notch signaling) (Fig. 3d). Collectively, these findings reveal a conserved, disease-enriched transcriptional state mainly defined by inflammatory and chemotactic gene programs.

### Proteomic profiling reveals a coordinated shift toward intracellular maintenance and away from membrane signaling

Comparable to transcriptomics, direct condition-level donor comparison revealed no genome-wide proteomic differences (Fig. S3). Filtering out donors with less than 3 fibers maintains no condition-level proteomic differences (Fig. S3). Extending the Inflammatory cluster assignment from the transcriptomic analysis, pseudobulk differential expression analysis on the proteomic dataset identified 277 differentially expressed proteins (DEPs), with 199 upregulated and 78 downregulated in cluster-assigned fibers (Fig. 4a). GO enrichment analysis of the 199 upregulated DEPs revealed strong enrichment for mitochondrial alterations, translational machinery, ribosomal biogenesis, and wound repair/matrix remodeling (Fig. 4b). The 78 downregulated DEPs showed enrichment for membrane-associated processes and surface protein localization (Fig. 4c). Notably, these proteomic annotations differ from the transcriptomic GO terms (immune activation, stress response), suggesting potential post-transcriptional or post-translational control mechanisms shaping the protein-level phenotype.

**Figure 4.**
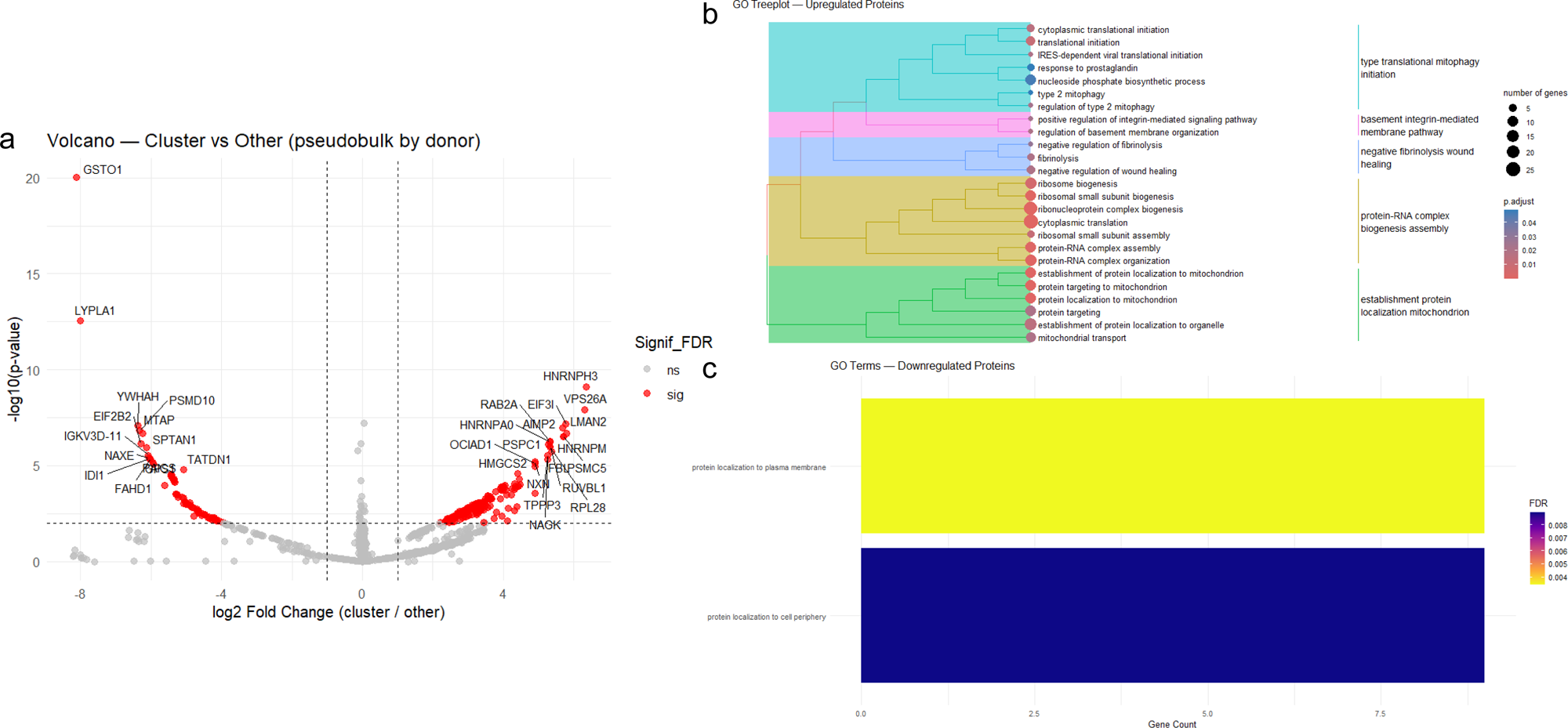
Cluster-Specific Proteomic Signatures in Single Myofibers. a) Volcano plot of DEPs between cluster and other. Fibers pseudobulked by donor and cluster status. Red dots indicate significant proteins. b) Upregulated GO terms for cluster DEPs. Treeplot with edges represents similarity between terms. c) Barplot of downregulated proteins, with fill showing FDR-adjusted p-value.

### RNA-protein integration identifies 19 concordantly altered features linking inflammation to intracellular remodeling

To assess concordance between transcriptomic and proteomic changes, the datasets were intersected based on shared gene symbols (Fig. S4). Directional agreement was assessed using fold changes; 19 features showed significant alterations in both datasets (Fig. 5, Table S2). Concordantly upregulated changes included ACTN4, MYL9, VTN, H2AC4, RPS9, CTSG, TPSB2, S100A11, CDSN. These proteins regulate cytoskeletal anchoring and dynamics (ACTN4, VTN, S100A11), myosin ATPase activity (MYL9), translational control (H2AC4, RPS9), and immune effector functions (CTSG, TPSB2, CDSN). Concordantly downregulated changes included only IDI1, involved in lipid metabolism and membrane biogenesis. Discordant regulations included AUH and NCK2; these showed decreased mRNA but increased protein levels, consistent with selective protein stabilization. Discordant regulations also included BOLA1, PRR33, SUN2, APOA2, FLOT2, IGKV4-1, and IGKV3D-11; these exhibited increased mRNA but decreased protein levels, suggesting enhanced degradation or secretion. Overall, these 19 overlapping features link the inflammatory transcriptional landscape (immune activation, cytotoxicity, ROS) to the proteomic shift toward mitochondrial and translational machinery, revealing a coordinated stress-adaptation program with selective post-translational control.

**Figure 5.**
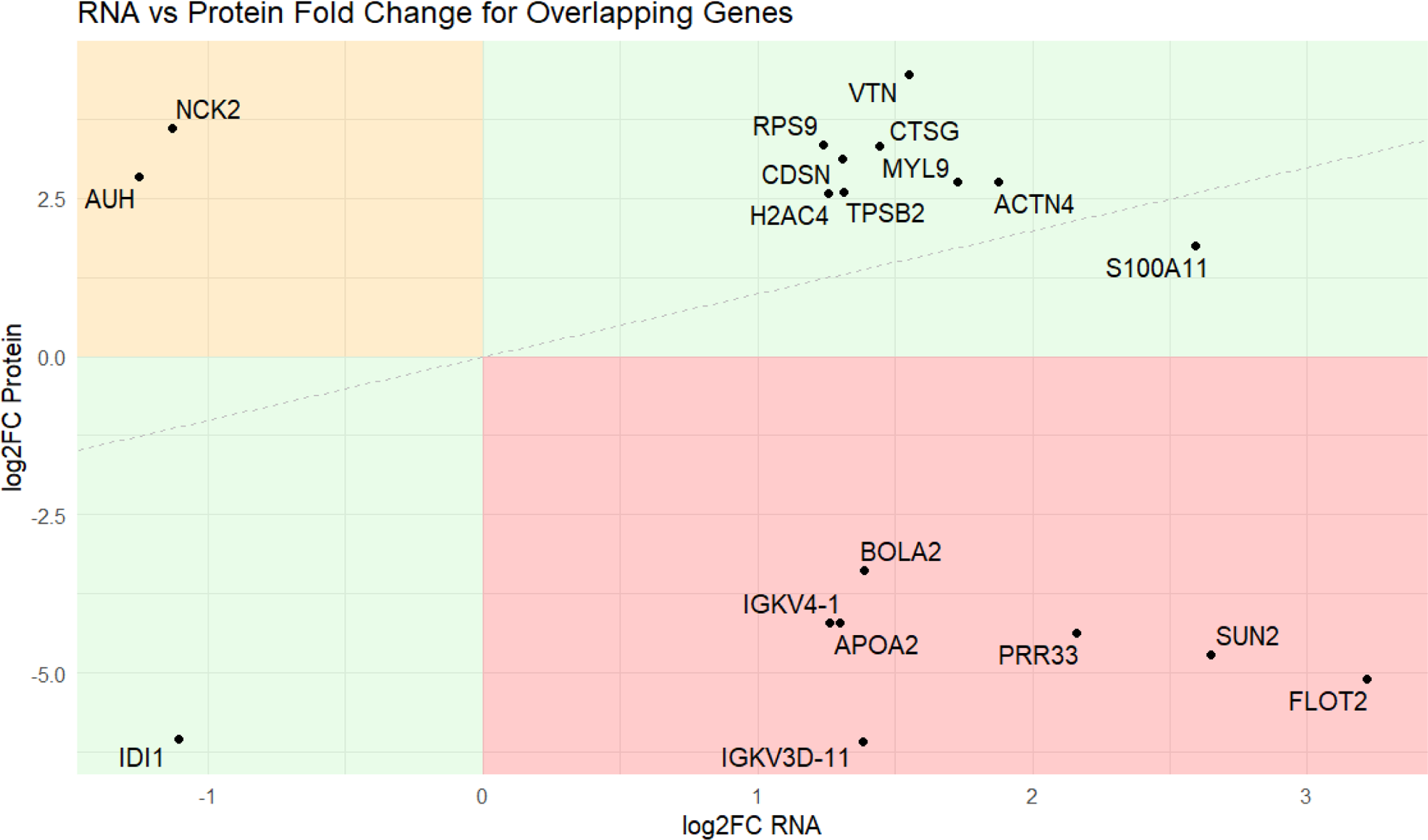
Concordance of RNA and protein fold changes for overlapping genes. Each point represents a feature significantly differentially expressed at both the transcript and protein level. Green = coordinate changes, orange = RNA down / protein up, red = RNA up / protein down. Dashed line represents a 1:1 relationship between RNA and protein fold changes.

### Inflammatory-clustered myofibers display prolonged ATP turnover time for the myosin super-relaxed state

To assess functional consequences of the Inflammatory cluster molecular phenotype, myosin ATP turnover kinetics were measured and correlated with the cluster-specific molecular features. For each fiber, a composite RNA–protein expression score was calculated as the average z-score-normalized expression across the 19 concordantly altered features. This score was tested for correlation with functional measurements: theoretical ATP consumption (Fig. 6a), DRX percentage (P1, Fig. 6b), SRX percentage (P2, Fig. 6c), DRX time constant (T1, Fig. 6d), and SRX time constant (T2, Fig. 6e).

**Figure 6.**
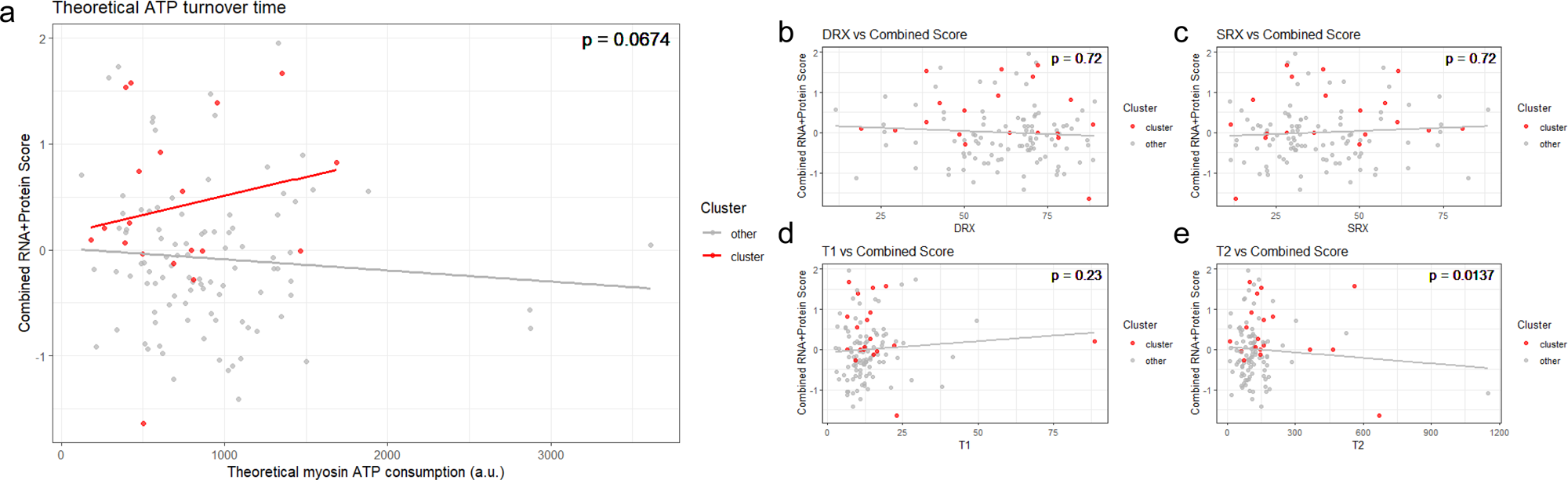
Relationship between functional fiber metrics and overlapping features. Per-fiber expression scores were computed with overlapping RNA–protein differentially expressed features. Scores were Z-transformed independently for RNA and protein and then averaged to create a score for each fiber. Wilcoxon rank-sum tests were performed to compare distribution of metric and average cluster/other score (p-value displayed per panel). Linear smooth is illustrative.

Fibers within the Inflammatory cluster showed significantly prolonged ATP turnover time in the SRX state (T2) compared to non-cluster fibers (Wilcoxon rank-sum test, p < 0.05). There was a significant difference in cluster composite score for T2 (p < 0.01), indicating that the molecular features defining the cluster are functionally linked to altered myosin energetics. Other kinetic parameters (T1, P1, P2, and overall ATP consumption) did not differ significantly between cluster and non-cluster fibers. This selective prolongation of SRX-state ATP turnover time indicates reduced basal ATPase activity in resting myosin molecules, consistent with bioenergetic adaptation to the inflammatory and stress-associated environment. The strong correlation between the multi-omic cluster score and functional measurements directly validates the molecular phenotype and establishes a link between transcriptomic/proteomic alterations and energy dynamics of the contractile apparatus, uniquely revealed with our workflow.

## Discussion

The complexity of the muscle microenvironment requires coordinated regulation across multiple cellular and molecular layers to maintain contractile function and metabolic homeostasis. Critical illness represents an acute systemic insult that disrupts this coordination, leading to rapid and profound muscle dysfunction known as ICU-AW (7, 8). Although numerous molecular profiling studies have characterized skeletal muscle changes in critical illness and ICU-AW (11, 12, 18–24), integration of multi-omic data with direct functional measurements at the level of individual cells remains limited. Building upon our recent developments (25), this study leverages a novel integrated approach that combines transcriptomic, proteomic, and functional myosin measurements from the same single myofiber to identify a disease-associated fiber state in ICU-AW. The key innovation is the direct cellular context in which molecular alterations are measured and validated through functional assessment, enabling interpretation of how coordinated programs impact contractile performance.

### A conserved disease-associated myofiber state emerges despite heterogeneity

Despite substantial heterogeneity in both ICU-AW and control populations (reflecting variation in age, sex, underlying conditions, and disease severity), a reproducible myofiber phenotype emerged within the ICU-AW cohort. This Inflammatory cluster phenotype was characterized by distinct transcriptional and proteomic signatures not present in control fibers. The conserved state is present in all ICU-AW donors and is not explained by myosin heavy chain composition. This suggests that critical illness induces a fiber-specific molecular program that, while not uniformly affecting all myofibers, represents a consistent tissue response to systemic stress.

The transcriptional profile of cluster-assigned fibers emphasized inflammatory and chemotactic processes (including immune activation, cytotoxicity, and granulocyte chemotaxis) indicating either a direct stress response or possible immune cell infiltration or interaction. Concurrently, the proteomic signature revealed a marked shift toward mitochondrial and translational machinery, with reduced membrane-associated signaling. This dissociation between transcriptomic and proteomic annotations suggests layered post-transcriptional and post-translational control.

### Multi-omic coordination reveals intracellular prioritization under stress

The identification of 19 concordantly altered features bridges the transcriptomic and proteomic layers to provide mechanistic insight. Firstly, a cytoskeletal and contractile remodeling is suggested by upregulation of ACTN4 and MYL9. This reflects altered actin anchoring and myosin ATPase modulation. ACTN4 overexpression destabilizes sarcomeres and induces a structural remodeling phenotype consistent with stress adaptation (26). Concurrent upregulation of VTN and S100A11 suggests enhanced cell adhesion and cytoskeletal plasticity (27, 28). Secondly, intracellular maintenance and mitochondrial changes can be linked to increased H2AC4 and RPS9. This indicates heightened translational activity, while downregulation of IDI1 reflects a shift away from lipid synthesis and membrane biogenesis, collectively supporting prioritization of intracellular repair machinery over extracellular communication. Thirdly, an immune effector integration may occur through elevated CTSG, TPSB2, and aberrant CDSN expression (a glycoprotein not normally expressed in skeletal muscle). This suggests either infiltrating immune cells or myofiber-intrinsic immune activation, possibly reflecting crosstalk with circulating cytokines or invading leukocytes. Fourthly and finally, our findings indicate a post-translational control through discordant features. AUH and NCK2 are increased despite reduced mRNA, suggesting selective protein stabilization. AUH dysregulation has been linked to mitochondrial defects and reduced stability of mature RNA transcripts (29), while NCK2 regulates actin cytoskeleton organization and cell migration (30). Discordance in elevated mRNA with reduced protein suggests selective degradation of BOLA1, PRR33, and SUN2. PRR33, in particular, is implicated in cytoskeleton–mitochondria organization, and its degradation reduces myofiber size and strength (31). This selective protein turnover suggests quality control and resource allocation mechanisms tuned toward mitochondrial maintenance and away from full-fiber growth and membrane-associated functions.

### Altered myosin super-relaxed state dynamics link molecular adaptation to energetic capacity

The functional finding of prolonged ATP turnover time for the SRX state connects the multi-omic phenotype to myosin energetics. The SRX state represents the resting conformation with minimal basal ATPase activity, and prolonged residence in this state reduces energetic cost during periods of inactivity. However, a shift toward SRX can also reflect impaired transition back to the active state, reducing maximal contractile capacity. The strong correlation between the composite multi-omic score and T2 (SRX time constant) validates the molecular cluster and suggests that stress-induced transcriptional and translational remodeling fundamentally alters myosin dynamics. This energetic shift aligns with the coordinated upregulation of MYL9 and ACTN4, known regulators of myosin ATPase activity and contractile dynamics. The proteomic emphasis on mitochondrial and translational machinery indicates a cell-autonomous bioenergetic adaptation (shifting resources toward maintenance and energy provision) at the expense of sustained contractile capacity. This may represent a compensatory cellular response to acute energy scarcity: by reducing basal ATP demand and prioritizing mitochondrial function, the stressed fiber sacrifices immediate contractile power for energetic survival.

### Methodological implications and biological significance

By integrating transcriptomic, proteomic, and functional measurements from individual myofibers, this work reveals cellular heterogeneity and functional adaptation beyond that which traditional bulk tissue analysis, or single-cell sequencing technologies, can capture. Single-fiber transcriptomics or proteomics alone might identify individual molecular changes; only the integrated approach reveals how these changes coordinate to reshape cellular function. This integrative framework is broadly applicable to complex disease states in which cellular heterogeneity obscures population-level signals. The single-fiber approach also mitigates a major confounding factor in bulk muscle studies: the mixture of fiber types, developmental states, and cell types within tissue. By measuring the same fiber across molecular and functional dimensions, the precision and interpretability of findings are substantially enhanced. This approach validates molecular signals within their native cellular context and reveals how adaptive programs couple to molecular and contractile layers.

### Clinical implications and therapeutic opportunities

The identification of a conserved disease-associated fiber state has important clinical implications. Muscle dysfunction in critical illness is a major determinant of morbidity and mortality, yet current therapeutic options are limited. The characterization of a fiber-specific molecular and functional phenotype opens new avenues for intervention. In finding a link between energetic shift and structural remodeling, this study supports a role for interventions that increase the availability of energy. Importantly, muscle dysfunction in critical illness may reflect a state of bioenergetic failure, in which impaired energy supply and/or utilization leads to early functional impairment that can precede overt structural damage. This framework further supports targeting mitochondrial dysfunction and inflammatory stress responses to preserve excitability and contractile performance (9). Furthermore, the multi-omic integration provides a framework for patient stratification: fibers or patients with higher abundance of the Inflammatory cluster phenotype might represent a distinct therapeutic target population.

### Limitations and future directions

Several limitations merit consideration. First, the sample size, while appropriate for a proof-of-concept, limits the bioinformatic models and approaches that could be used. Second, this study measures ATP turnover kinetics but does not directly assess contractility or force generation; future work incorporating force measurements would clarify the functional impact of altered myosin dynamics on contractile performance. Third, the identity of cells expressing immune-associated genes (infiltrating immune cells versus intrinsic myofiber expression) remains to be determined; single-nucleus RNA-seq or immunofluorescence would resolve this question.

Future studies incorporating larger cohorts will be essential for validating the conserved fiber state and establishing its prevalence and clinical relevance across different ICU populations and disease etiologies. Extended proteomic analyses could reveal post-translational modifications (phosphorylation, acetylation, ubiquitination) that underlie the selective protein stabilization and degradation observed here. Finally, mechanistic studies in *in vitro* systems or animal models could test whether targeting the identified molecular features restores myosin dynamics and contractile capacity under stress conditions.

### Conclusions

This study suggests that in critically-ill patients, ICU-AW induces a conserved, fiber-specific molecular and functional phenotype characterized by inflammatory signaling, intracellular remodeling, and altered myosin energetics. By directly linking multi-omic alterations to myosin dynamics at the level of individual myofibers, this work provides a broadly applicable integrative framework for dissecting cellular heterogeneity and functional adaptation in complex human diseases. The identification of coordinated transcriptomic and proteomic programs, validated through functional measurement, suggests that targeting this fiber state may offer new therapeutic opportunities in ICU-acquired dysfunction and related muscle disorders.

## Methods

### Sex as a biological variable

ICU-AW affects both males and females. For this reason, the present study includes a mixture of sexes.

### Study approval

Here, investigations only included human tissue. Note that (i) all clinical information and materials were obtained for diagnostic purposes; (ii) informed consent was obtained for the patient or legal representative; (iii) procedures conformed to the standards set by the latest version of the Declaration of Helsinki and Good Clinical Practice; and (iv) the study was approved by the Ethics Committee of Brescia (Protocol number 3468). Additional biopsies were collected in the context of a single-blinded, randomized controlled trial (RCT), which was discontinued due to low inclusion rates. The protocol has been filed on 27 July 2017 in the Clinical Trial Register under #NCT03231540 and was approved by the Medical Ethical Committee of VU Medical Center, Amsterdam, The Netherlands.

### Human Participants, Tissue Collection, and Storage

Skeletal muscle tissue biopsies were obtained from 16 subjects hospitalized in the Intensive Care Unit (ICU-AW group, n=8) and Orthopaedic ward (control donors, n=8) of the Spedali Civili University Hospital in Brescia, Italy and from VU Medical Center, Amsterdam, Netherlands. All traceable identifiers were removed before analysis to protect patient confidentiality; all samples were analyzed anonymously. Informed consent was obtained from the patient before the collection of samples. The subjects were both male and females, with an age range of 19-82 years (average age 62 y/o).

Human skeletal muscle tissue was dissected from quadriceps femoris. The muscle samples harvested from patients hospitalized in ICU were obtained with a surgical biopsy, 8-10 days after ICU admission. The skin was prepped with antiseptic Clorhexidin 2% solution and draped. A longitudinal 4 cm midline incision was performed at the middle third of the thigh. The quadriceps muscular fascia was reached through the subcutaneous tissue and longitudinally incised. Hemostasis was obtained in every case by compression. The quadriceps fascia and the subcutaneous tissue were sutured with 3-0 polyglactin 910 braided adsorbable suture; the skin was sutured with 4-0 polyglactin 910 braided adsorbable suture. A compressive dressing was applied at the end of the procedure. The samples obtained from patients in the Orthopedic Ward were obtained during total hip replacement performed for hip arthritis through a direct anterior approach. As soon as the biopsy was removed from the patient, it was immediately placed into a polypropylene tube filled with 25 ml cold (+4°C) NaCl 0.9% solution and washed twice. Skeletal muscle tissue was, then, moved to a new tube and snap-frozen in liquid nitrogen. Samples was stored at -80°C.

### Single Myofiber Isolation and Preparation

On the day of functional profiling, muscle samples were thawed briefly and transferred to cold relaxing buffer (composition: 4 mM Mg-ATP, 1 mM free Mg²⁺, 10⁻^9^ µM free Ca²⁺, 20 mM imidazole, 7 mM EGTA, 14.5 mM creatine phosphate, and KCl to adjust ionic strength to 180 mM; pH 7.0). Working rapidly to minimize RNA and protein degradation, individual myofibers were manually isolated under a dissecting microscope.

Individual myofibers were positioned on custom-prepared microscopy slides fitted with half-split copper transmission electron microscopy grids. Clamped myofibers were maintained on ice during isolation. Once all myofibers were clamped, the unclamped tail segments were carefully excised and placed into microtubes containing RLT+ buffer. The spatial location of each fiber tail was meticulously recorded for later integration with functional data. Lysates were stored at −80°C pending RNA and protein isolation.

### Mant-ATP Chase Experiments for Myosin ATP Turnover Kinetics

Clamped myofibers on slides were permeabilized with 0.1% Triton X-100 in relaxing buffer for 10 minutes at room temperature. A coverslip was then positioned over the grid and myofibers to form a flow chamber. The slide was incubated for 5 minutes in rigor buffer (composition: 120 mM potassium acetate, 5 mM magnesium acetate, 2.5 mM K₂HPO₄, 50 mM MOPS, 2 mM DTT; pH 6.8). The solution was then exchanged for rigor buffer containing 250 μM Mant-ATP and incubated in darkness for an additional 5 minutes to label myosin heads.

After labeling, the slide was rapidly flushed with rigor buffer containing 4 mM ATP to initiate the ATP chase. Simultaneously, fluorescence decay was captured using an inverted Zeiss Axio Observer microscope. Two reference images were acquired immediately before ATP addition. Subsequent images were recorded every 5 seconds for a total of 5 minutes.

Fluorescence intensity within individual myofibers was analyzed using ImageJ. For each frame, mean background fluorescence intensity was subtracted from the fiber-specific fluorescence intensity. All intensities were normalized to the final Mant-ATP image before washout.

The normalized fluorescence decay curves were fit to an unconstrained double exponential model:

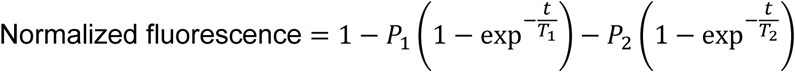

In this model: P₁ = amplitude of the initial rapid decay (disordered-relaxed myosin heads); T₁ = time constant of the fast decay phase (DRX ATP turnover rate); P₂ = amplitude of the slower second decay (super-relaxed myosin heads); T₂ = time constant of the slow decay phase (SRX ATP turnover rate); Theoretical ATP consumption was solely calculated from these parameters (32).

### RNA Isolation and Sequencing Library Preparation

mRNA was captured from 10 μL of fiber lysate using the NEBNext Poly(A) mRNA Magnetic Isolation Module according to the manufacturer’s instructions. The resulting supernatant was retained for protein precipitation.

Captured mRNA was converted into Illumina-compatible cDNA libraries using the NEBNext Single Cell/Low Input RNA Library Prep Kit. Reverse transcription and template switching were performed, followed by PCR amplification for 14 cycles. Library quality was assessed on an Agilent TapeStation 4200, and concentration was determined using a Thermo Fisher QubitFlex fluorometer.

Per sample, 2 ng of cDNA was fragmented, end-repaired, and ligated with Illumina adapters. Libraries were amplified for an additional 8 PCR cycles with unique NEBNext Multiplex Oligos. Fragment size and molarity were assessed on the TapeStation. Libraries were pooled at equimolar concentration (2 nM) and sequenced on a NovaSeq X Plus Series platform to generate 150 bp paired-end reads.

### Primary RNA-Seq Data Processing

Raw RNA sequencing data were quality-controlled and aligned using the nf-core/rnaseq pipeline (version 3.19.0). Prior to pipeline execution, reads were trimmed using Flexbar according to recommendations from New England Biolabs. Alignment was performed with STAR, and transcript abundance was quantified with Salmon. Output count matrices were used for downstream analysis.

### Sample preparation for Proteomics Analyses

The supernatant from mRNA isolation was retained for protein analysis, performed as previously described (25). Ice-cold 100% acetone in 100 mM Tris-HCl (pH 8.5) was incubated with proteins overnight. Precipitation was performed with sequential centrifugation and washing in 100% acetone/100 mM Tris buffer. Acetone was evaporated from the samples using a SpeedVac concentrator. Pellets were resuspended in 10 μL of digestion buffer (composition: 1% sodium deoxycholate, 10 mM DTT, 40 mM chloroacetamide, 50 mM Tris-HCl pH 8.5) and boiled at 95°C with shaking for 10 minutes. Samples were briefly centrifuged, then sonicated for 15 minutes in a water bath sonicator.

Protein digestion was performed overnight at 37°C using a mix of trypsin and Lys-C (1:100 and 1:500 enzyme to protein ratios, respectively). The next day, peptides were desalted using in-house-prepared Stage-Tips packed with Styrene divinylbenzene-reverse phase sulfonate (SDB-RPS) disks.

Digested samples were transferred onto tips and centrifuged until complete loading. Tips were first washed with 1% trifluoroacetic acid (TFA) in isopropanol followed by a wash with 0.2% TFA in water. The resulting desalted peptides were eluted with 80% acetonitrile and 1% ammonia solution. Samples were dried in a SpeedVac and resuspended in A* buffer (5% acetonitrile, 0.1% TFA). Samples were transferred to Evotips according to the manufacturer’s instructions prior to LC-MS/MS analysis.

### LC-MS/MS

LC–MS/MS analyses were performed as previously described (25). The LC instrumentation consisted of an Evosep One HPLC system (Evosep, Odense, Denmark) coupled by electrospray ionisation to a timsTOF HT SCP mass spectrometer (Bruker, MA, USA). Peptide separation was achieved on an 8 cm column (150 µm inner diameter) packed with 1.5 µm C18 particles. Chromatography was conducted using the “60 samples per day” gradient method, with ionisation via a CaptiveSpray2 source and a 20 µm emitter for MS introduction. Single muscle fibre peptide samples were analysed using DIA-PASEF, employing an acquisition of 100-1700 m/z range. The shot-gradient diaPASEF method was used, including 8 diaPASEF scans with three 25 Da windows per ramp, a total mass range of 475-1000 Da, and a mobility range of 1.30-0.85 1/K0. The collision energy was decreased linearly from 59 eV at 1/K0 = 1.6 to 20 eV at 1/K0 = 0.6 Vs cm-2. Both accumulation time and PASEF ramp time were set to 100 ms, with a total cycle time of 0.95 s across 8 DIA-PASEF scans.

### Primary MS data processing

MS spectra were processed with DIA-NN software (version 2.0) (33), using a myofiber-specific library developed in-house (4). Protein group quantification was based exclusively on proteotypic peptides, with the neural network configured to operate in double-pass mode. The quantification strategy was set to “Robust LC (high accuracy),” and match-between-runs was activated. All other parameters were left at default values, including a peptide length window of 7–30 amino acids and a precursor-level FDR cutoff of 1%. Downstream bioinformatics were conducted using the PG_matrix file from the DIA-NN output.

### Transcriptomic Analysis

All bioinformatics analyses were performed in R (version 4.4.2). Raw transcriptomic count matrices and sample metadata were converted into a Seurat object (version 5.3.0) (34) for downstream analysis and then converted to a Single Cell Experiment (version 1.28.0) object for quality control and normalization.

Quality control filtering removed fibers with gene counts or feature abundance ±3 median absolute deviation (MAD) from the median. Fibers meeting QC criteria were retained for subsequent analysis (final n = 165 fibers). Library-size normalization was performed using LogNormcounts (Scran version 1.34.0)(35). Batch effects were corrected using Harmony (36).

### Differential Expression Analysis

Differential expression analysis was conducted using a pseudobulk approach to account for the nested donor structure. Raw counts were aggregated across all fibers within each donor-condition combination. Pseudobulk counts were normalized using the trimmed mean of M-values (TMM, edgeR version 4.4.2) (37) method and analyzed using limma–voom (version 3.62.2) (38). Count data were converted to log₂ counts per million (CPM), and linear regression modeling with empirical Bayes moderation was applied.

Differentially expressed genes were ranked by false discovery rate (FDR) using the Benjamini-Hochberg method; significance threshold was FDR < 0.05.

### Gene Set Enrichment Analysis

GSEA was performed on the ranked DEG list using the fgsea algorithm (version 1.32.4) (39) with 10,000 permutations. Hallmark gene sets from MSigDB (version 25.1.1) (40) were tested for enrichment. Normalized enrichment scores (NES) and FDR-adjusted p-values were calculated.

### Cluster Identification and Characterization

The top 50 DEGs were selected for visualization at the single-fiber level. Expression values were z-score normalized across all fibers. Hierarchical clustering (Euclidean distance, complete linkage) and heatmap visualization were performed using pheatmap (version 1.0.13) (41). Fibers were assigned to cluster or non-cluster groups based on dendrogram membership.

To characterize the cluster phenotype, differential expression analysis compared cluster-assigned versus non-cluster fibers, aggregated by donor and cluster status.

Overexpression gene Ontology (GO) enrichment analysis was performed using the clusterProfiler package (version 4.14.6) (42) separately for up- and downregulated genes.

### Proteomic Processing and Analysis

Proteomic data (protein intensity matrix) were log₂-transformed. Samples with more than 50% missingness were filtered out. Proteins were retained if quantified in at least 70% of samples within each condition. Between-sample normalization was performed using the scale method in limma. Missing values were imputed using the tImpute() function from the PhosR package (version 1.16.0) (43).

Fiber types were assigned based on the relative abundance of myosin heavy chain (MYH) isoforms measured at the protein level. Log2-normalized and imputed protein intensities for MYH7, MYH2, and MYH1 were extracted for each individual fiber and transformed back to linear scale. For each fiber, the relative contribution of each MYH isoform was calculated as a percentage of the total MYH signal (MYH7 + MYH2 + MYH1). Fibers were then ordered from highest to lowest relative expression for each MYH isoform independently. Data-driven thresholds for dominant isoform expression were determined using a bottom-knee detection approach based on the point of maximal curvature in the ordered distribution. Fibers not meeting criteria for a dominant or hybrid classification were labeled as other hybrid fibers (4).

Differential protein expression analysis was performed using the same pseudobulk approach as the transcriptomic analysis. GO enrichment analysis was conducted separately for up- and downregulated proteins.

### Multi-Omic Integration

DEGs and DEPs were intersected based on shared gene symbols. Directional agreement was assessed by comparing the signs of fold changes. Overlapping features (n = 19) were extracted from the significantly changed RNA and protein expression matrices. For each fiber, average expression values across the 19 overlapping features were computed separately for RNA and protein data. These averages were z-score normalized and averaged to generate a single composite RNA–protein score per fiber.

### Functional Correlation Analysis

Associations between the composite score and functional measurements (P1, T1, P2, T2, and theoretical ATP turnover time) were tested using non-parametric Wilcoxon rank-sum tests. Linear regression was used to estimate the relationship between the composite score and functional parameters.

### Quantification and Statistical Analysis

All reported p-values for differential expression are FDR-adjusted. FDR was controlled using the Benjamini-Hochberg method. Differential expression: FDR < 0.05 was used as the significance threshold for DEGs and DEPs. Pathway enrichment analyses used FDR adjustment. Functional associations: Associations between the composite score and functional measurements were tested using Wilcoxon rank-sum tests with two-tailed p-values. Significance threshold: p < 0.05.

Sample sizes: Transcriptomic analysis included 165 individual fibers (control n = 83, ICU-AW n = 82) from 16 donors. Proteomic analysis included 124 fibers (control n = 62, ICU-AW n = 62) from 16 donors. Cluster-versus-non-cluster analysis included 117 fibers with both datasets (ICU-AW-cluster n = 32, non-cluster ICU-AW n = 17, control non-cluster n = 68).

### Availability of Data and Material

All the data analyzed are presented here in the figures, supplementary files and public data repositories. Specifically, RNA-sequencing data have been deposited in the European Genome-phenome Archive under study ID [submitted]. The mass spectrometry proteomics data have been deposited to the ProteomeXchange Consortium via the PRIDE partner repository with the dataset identifier PXD073993. All bioinformatics code and analysis scripts are available on GitHub (https://github.com/amwinant/ICUAW-SFMO-pilot.git). Custom R scripts for data processing, analysis, visualization, and statistical testing are available on GitHub. The repository includes data loading and QC filtering, pseudobulk differential expression pipelines, GSEA and pathway analysis, GO enrichment, hierarchical clustering and cluster assignment, multi-omic integration and score calculation, functional correlation analysis, and figure generation.

## Author contributions

AMW, LP, SP, NL, RAES and JO contributed to the study conception and design. Material preparation, data collection and analysis were performed by AMW, RMJ, LP, WJC, CACO, ASD, SC, SP, NL, and RAES. The first draft of the manuscript was written by AMW and JO and all authors commented on all versions of the manuscript and approved the submitted version.

## Acknowledgments

We thank Thomas Nyegaard Beck for his laboratory support. This work was generously funded by the Lundbeckfonden (grant agreement number R434-2023-311) to JO. Additionally, mass spectrometry analyses were performed by the Proteomics Research Infrastructure (PRI) at the University of Copenhagen (UCPH), supported by the Novo Nordisk Foundation (grant agreement number NNF19SA0059305). RS is supported by the Medical Research Council (UKRI2537) and the Acadmey of Medical Sciences (SBF0010\1048). The original trial for which some of the biopsies were collected was supported by an unrestricted grant from Nestlé. Nestlé had no role in the design of the study and collection, analysis and interpretation of the data or in writing the manuscript. CO was supported by NHLBI Grant HL-121500. This work was further supported by the Italian Ministry of Health (Ministero della Salute) under the Ricerca Finalizzata (RF) “Theory enhancing” grant: ID project RF-2021-12375279 and “Alessandra Bono” Foundation.

