## Supplemental Information for "Integrated Single-Fiber Multi-Omics Links an Inflammatory-Associated Myofiber State to Altered Myosin Dynamics in Patients with ICU-acquired weakness"

This document includes:

- Tables S1 and S2
- Figures S1 – S4

**Table S1. Workflow and final fiber inclusion information.**

| Donor ID | Condition | Fibers in functional analysis | Fibers in transcriptomic analysis | Fibers in proteomic analysis | Overlapping fibers for integration analysis |
|---|---|---|---|---|---|
| **1** | Control | 12 | 11 | 10 | 10 |
| **2** | Control | 12 | 11 | 10 | 10 |
| **3** | Control | 12 | 12 | 6 | 6 |
| **4** | Control | 12 | 10 | 10 | 8 |
| **5** | Control | 12 | 19 | 10 | 10 |
| **6** | Control | 12 | 11 | 7 | 6 |
| **7** | Control | 12 | 8 | 7 | 6 |
| **8** | Control | 12 | 9 | 7 | 7 |
| **1** | ICU-AW | 12 | 8 | 1 | 1 |
| **2** | ICU-AW | 12 | 8 | 3 | 2 |
| **3** | ICU-AW | 12 | 10 | 9 | 8 |
| **4** | ICU-AW | 12 | 11 | 11 | 11 |
| **5** | ICU-AW | 12 | 12 | 11 | 11 |
| **6** | ICU-AW | 12 | 11 | 10 | 9 |
| **7** | ICU-AW | 12 | 11 | 7 | 7 |
| **8** | ICU-AW | 12 | 11 | 5 | 5 |

**Table S2. Gene annotations for overlapping RNA–protein differentially expressed features.** Manually annotated functional descriptions for features exhibiting concordant differential regulation at both the transcriptomic and proteomic level. Annotations were compiled from NCBI gene summaries.

| Gene | Full name | Function summary |
|---|---|---|
| **ACTN4** | Actinin Alpha 4 | Helps anchor the myofibrillar actin filaments |
| **MYL9** | Myosin light chain 9 | Regulates muscle contraction by modulating the ATPase activity of myosin heads |
| **IDI1** | Isopentenyl-diphosphate delta isomerase 1 | Peroxisomally-located, catalyzes substrate for ultimate synthesis of cholesterol |
| **VTN** | Vitronectin | Promotes cell adhesion, migration, ECM degradation |
| **H2AC4** | H2A clustered histone 4 | Replication-dependent histone that is a member of the histone H2A family |
| **RPS9** | Ribosomal protein S9 | Ribosomal protein that is a component of the 40S subunit |
| **CTSG** | Cathespin G | Component of azurophil granules of neutrophilic polymorphonuclear leukocytes |
| **TPSB2** | Tryptase beta 2 | Trypsin-like serine protease, main isoenzymes expressed in mast cells |
| **CDSN** | Corneodesmosin | Glycoprotein typically localized to human epidermis |
| **S100A11** | S100 calcium binding protein A11 | Motility, invasion, and tubulin polymerization |
| **AUH** | AU RNA binding methylglutaconyl-CoA hydratase | Involved in RNA degradation and deadenylation, mitochondrial protein synthesis |
| **NCK2** | NCK adaptor protein 2 | Cytoskeletal reorganization |
| **BOLA1** | bolA family member 1 | Cell redox homeostasis, mitochondrial protein |
| **IGKV4-1** | Immunoglobulin kappa variable 4-1 | Antigen binding activity, immune response |
| **APOA2** | Apolipoprotein A2 | Second most abundant protein of the high density lipoprotein particles |
| **PRR33** | Proline rich 33 | Wound healing |
| **SUN2** | Sad1 and UNC84 domain containing 2 | Inner nuclear membrane, connection between nuclear lamina and cytoskeleton |
| **FLOT2** | Flotillin 2 | Vesicular trafficking and signal transduction |
| **IGKV3D-11** | Immunoglobulin kappa variable 3D-11 | Immune response, located in extracellular region and plasma membrane |

**Figure S1. Jaccard similarity heatmap showing pairwise gene set overlap among MSigDB Hallmark pathways.** The Jaccard index reflects the proportion of shared genes between two pathways relative to the size of their union.

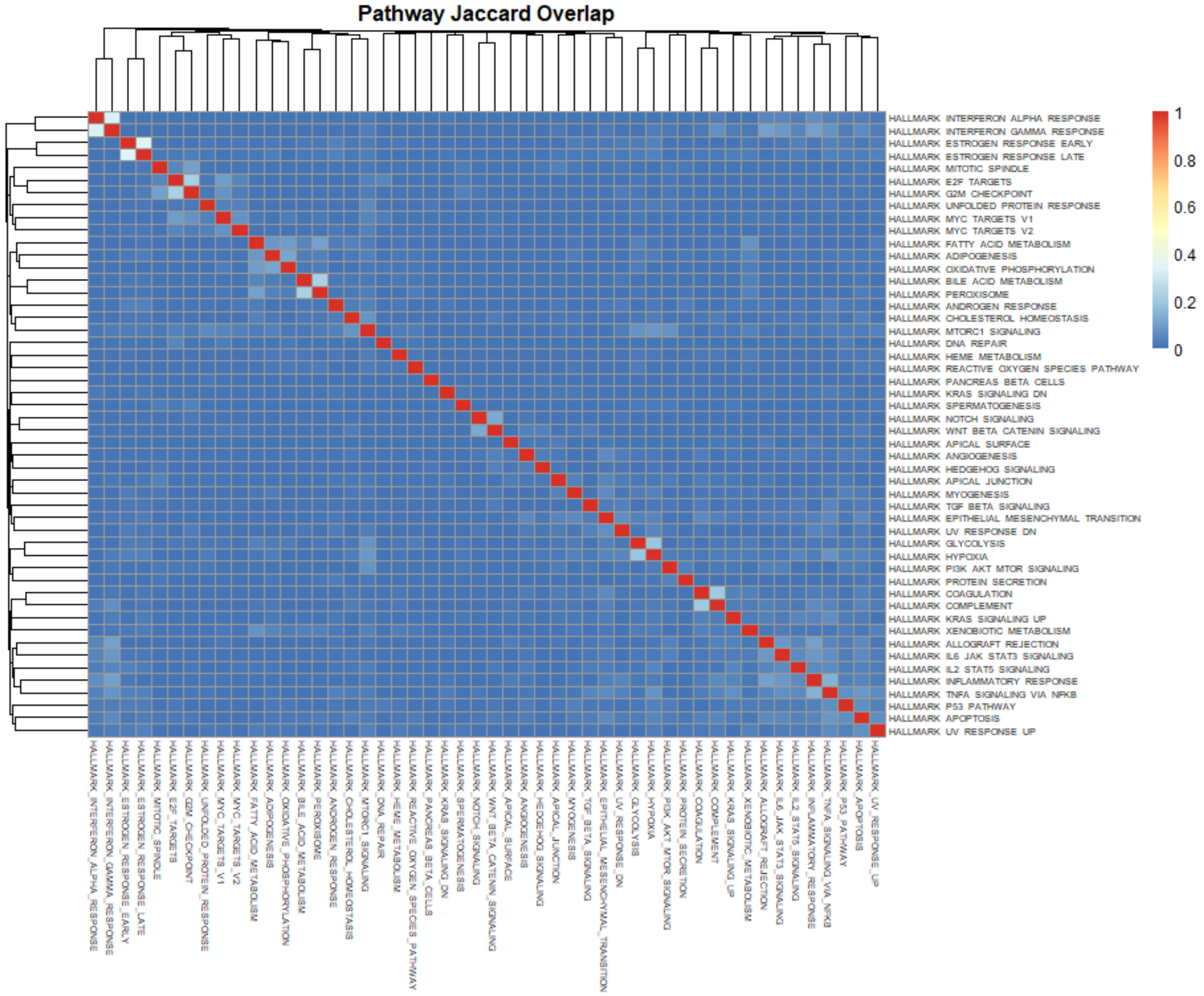

**Figure S2. Cluster composition.** a) Stacked barplot colored by fiber type (based on myosin heavy chain expression), showing composition of cluster group compared to other fibers. b) Feature counts per fiber, showing comparable coverage of cluster fibers to other fibers.

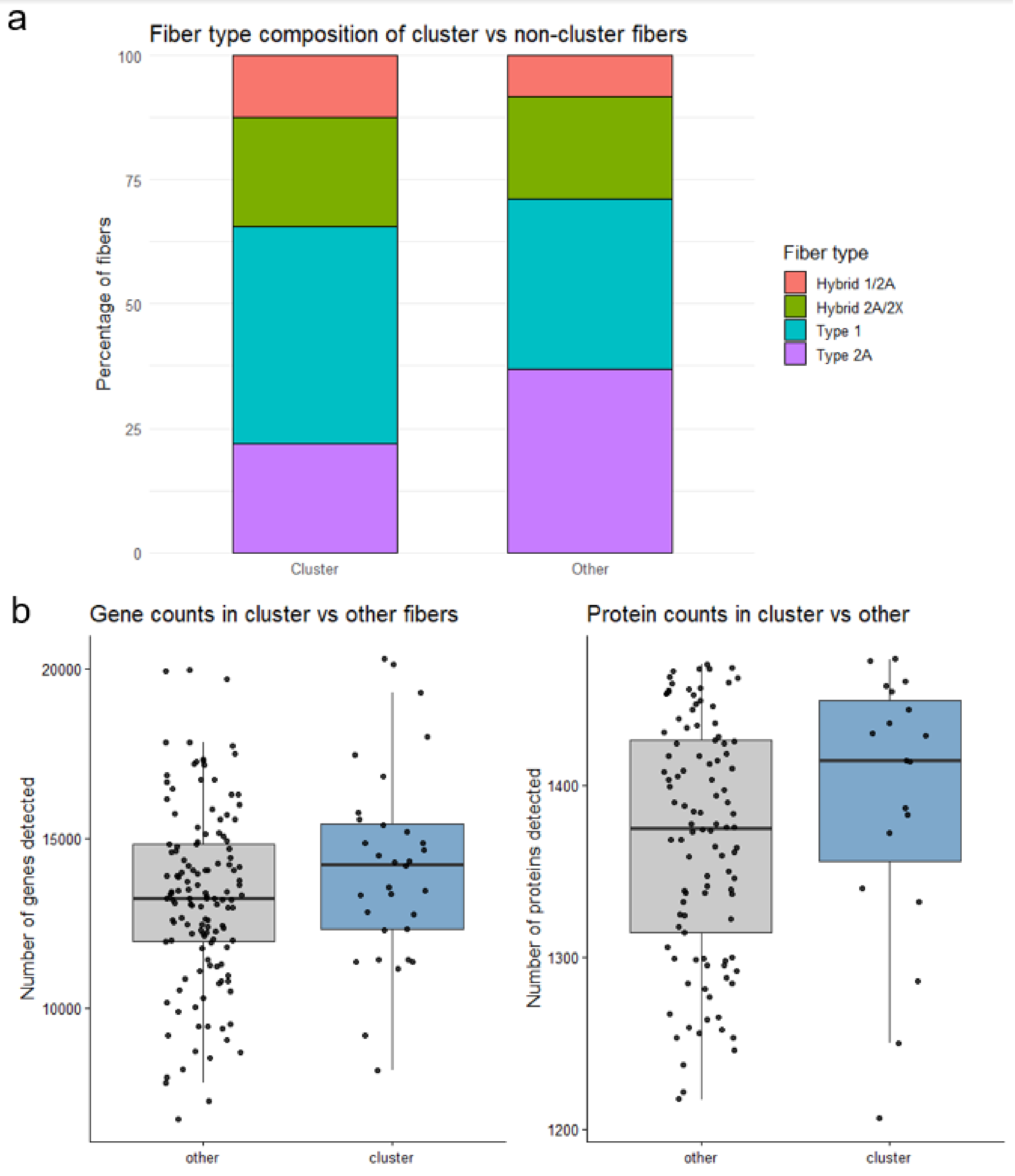

**Figure S3. DEPs between ICU-AW and control samples.** a) Volcano plot with fibers aggregated by donor. Gray dots indicate non-significant proteins. b) Number of fibers per donor; donors with less than 3 fibers are highlighted. c) Volcano plot with fibers aggregated for donors with at least 3 fibers. Gray dots indicate non-significant proteins.

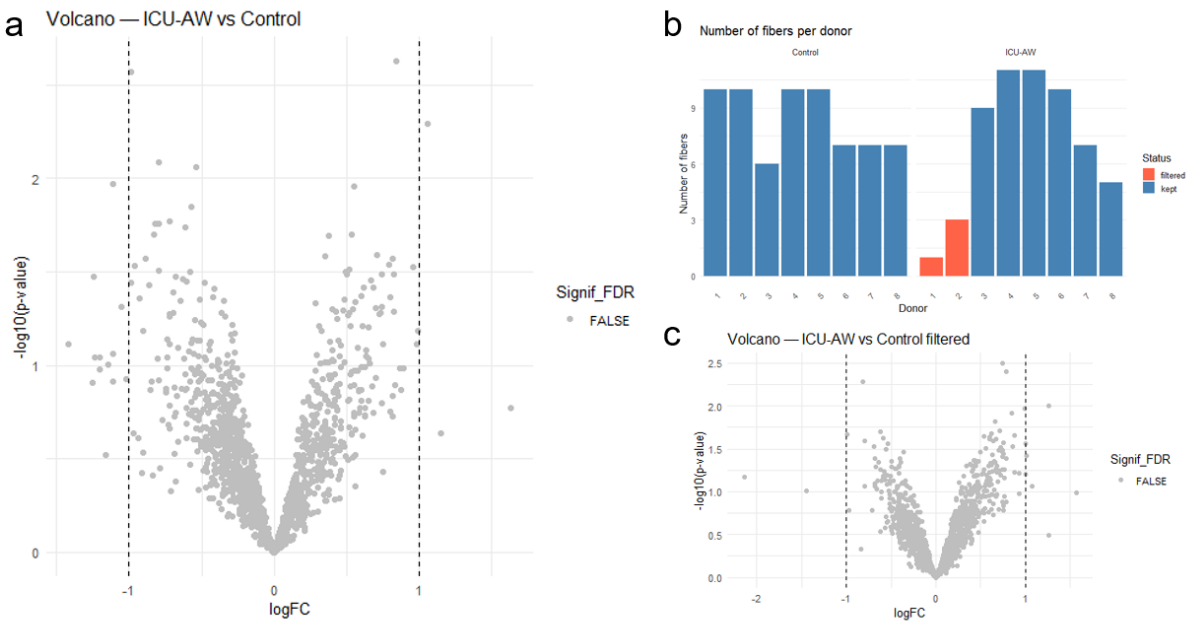

**Figure S4. Concordance of RNA and protein fold changes for all features.** Each point represents the fold changes of a feature differentially expressed at both the transcript and protein level. Red points are features significantly changed in both RNA and protein datasets. Dashed line represents a 1:1 relationship between RNA and protein fold changes.

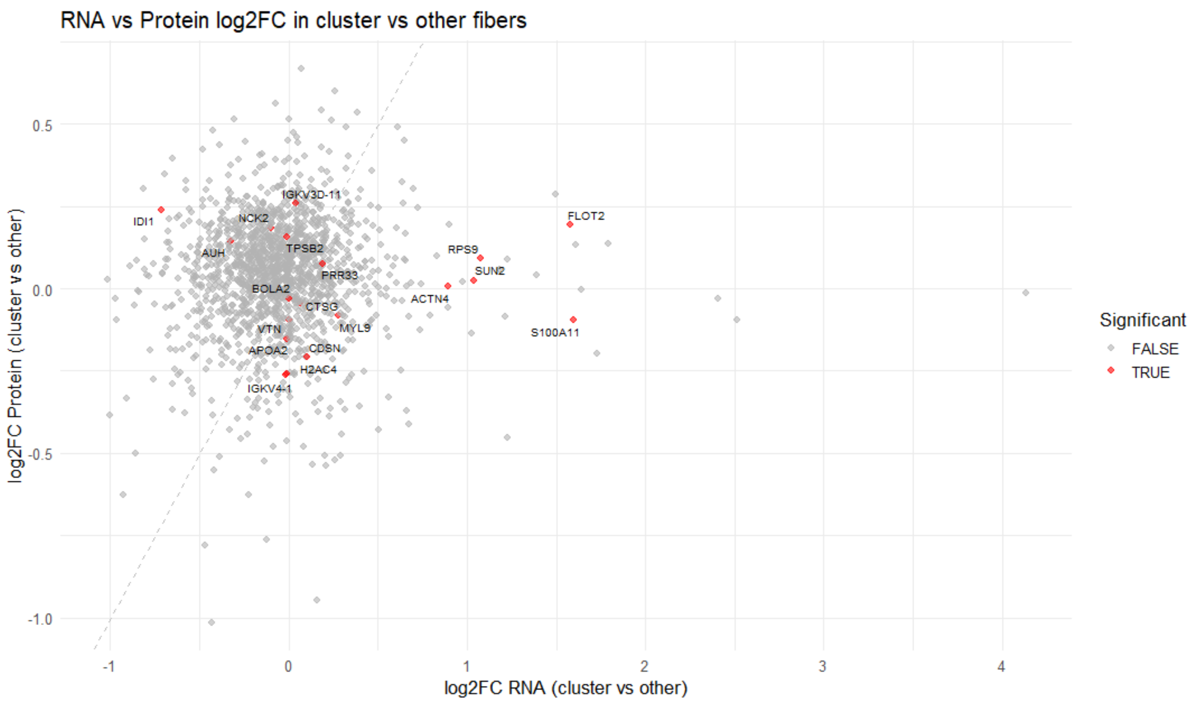
